# PRAK-03202: A triple antigen VLP vaccine candidate against SARS CoV-2

**DOI:** 10.1101/2020.10.30.360115

**Authors:** Saumyabrata Mazumder, Ruchir Rastogi, Avinash Undale, Kajal Arora, Nupur Mehrotra Arora, Biswa Pratim, Dilip Kumar, Abyson Joseph, Bhupesh Mali, Vidya Bhushan Arya, Sriganesh Kalyanaraman, Abhishek Mukherjee, Aditi Gupta, Swaroop Potdar, Sourav Singha Roy, Deepak Parashar, Jeny Paliwal, Sudhir Kumar Singh, Aelia Naqvi, Apoorva Srivastava, Manglesh Kumar Singh, Devanand Kumar, Sarthi Bansal, Satabdi Rautray, Indrajeet Singh, Pankaj Fengade, Bibekanand Kumar, Manish Saini, Kshipra Jain, Reeshu Gupta, Prabuddha Kumar Kundu

**Affiliations:** Premas Biotech Private Limited, Manesar, India

## Abstract

The rapid development of safe and effective vaccines against SARS CoV-2 is the need of the hour for the coronavirus outbreak. Here, we have developed PRAK-03202, the world’s first triple antigen VLP vaccine candidate in a highly characterized *S. cerevisiae-based* D-Crypt^TM^ platform, which induced SARS CoV-2 specific neutralizing antibodies in BALB/c mice. Immunizations using three different doses of PRAK-03202 induces antigen specific (Spike, envelope and membrane proteins) humoral response and neutralizing potential. PBMCs from convalescent patients, when exposed to PRAK-03202, showed lymphocyte proliferation and elevated IFN-γ levels suggestive of conservation of epitopes and induction of T helper 1 (Th1)–biased cellular immune responses. These data support the clinical development and testing of PRAK-03202 for use in humans.

## Main

The outbreak of Coronavirus Disease 2019 (COVID-19) was declared a Public Health Emergency of International Concern on 30 January 2020, and a pandemic on 11 March 2020 by WHO. As of September-2020, with more than ~864,000 deaths and ~250,000 new cases each day, it is considered a serious threat to global public health. There are currently no effective therapeutics^1^, calling for the urgent development of safe and effective therapeutics against nCOVID19.

SARS CoV-2 is an enveloped virus with a positive-strand RNA of 29.9 kilobases and contains four structural proteins, including spike (S), envelope (E), membrane (M), and nucleocapsid (N) proteins. Currently, most efforts have focussed on developing DNA and RNA vaccines to combat COVID-19. However, there are several challenges including, I) Poor immunogenicity II) single target-oriented approach for the current clinical trial vaccines-RBD or spike protein III) amounts and duration of protein production, and IV) chances of integrating foreign DNA into the host genome2 V) high mutation rate of spike protein, VI) ultra-cold chain storage of the vaccines, thus creating a bottleneck in the development of an effective COVID-19 vaccine. While spike protein has been well documented as a target for developing effective vaccines or drug-mediated therapies, recent studies showed the significance of M and E proteins in efficient assembly, release, and secretion of SARS CoV-2^1,3^. Moreover, competition between neutralizing and non-neutralizing epitopes of spike protein could greatly reduce the host immune response^3^ and increased failure chances of single-antigen targeting vaccines.

Immunization with a virus like particles (VLPs) has been shown to be an effective strategy for vaccine development for respiratory viruses by several groups as they appear to be improved in terms of biological safety^4–7^. This study aims to establish approaches to overcome the abovementioned limitations by producing a VLP based vaccine candidate, PRAK-03202 for COVID19.

In this study, we developed a VLP based COVID-19 vaccine, PRAK-03202, to defeat the nCOVID-19 virus. PRAK-03202 design and synthesis were initiated upon the public release of the SARS-CoV2 genome sequences in February 2020. Co-expression of Spike (S), Envelope (E), and Membrane (M) proteins (SEM) have been shown to form VLPs for SARS-CoV8. The design and manufacture of the PRAK-03202 vaccine candidate represent a plug and play process in which we insert the three-target antigen (S, E, and M) sequence into a highly characterized *S.cerevisiae* based D-Crypt^TM^ platform (Premas Biotech). This system allows the drug product’s scalability, and thus circumventing conventional vaccine production complexities in eggs or cell culture^9^. PRAK-03202 was purified from the cell lysates and analysed for expression of SEM proteins. Immunoblot analysis confirmed the expression of the SEM proteins using commercially available antibodies (Figure 1A). The co-expression of S, E and M was further confirmed by mass spectrometry. The purified PRAK-03202 showed a purity of greater than 98% using SEC-HPLC (Figure 1B). PRAK-03202 was further characterized by DLS (Dynamic Light Scattering) and Cryo-Transmission Electron Microscopy (Cryo-TEM). DLS showed a mean particle size of 176 ± 18 nm with 18% polydispersity index (Figure 1C-). Additionally, cryo-electron microscopy (cryo-EM) analysis showed intact spherical-shaped bilayer particles^10^ with distinct crown-like spikes representing a prefusion state of the virus (Figure 1D).

**Figure 1.**
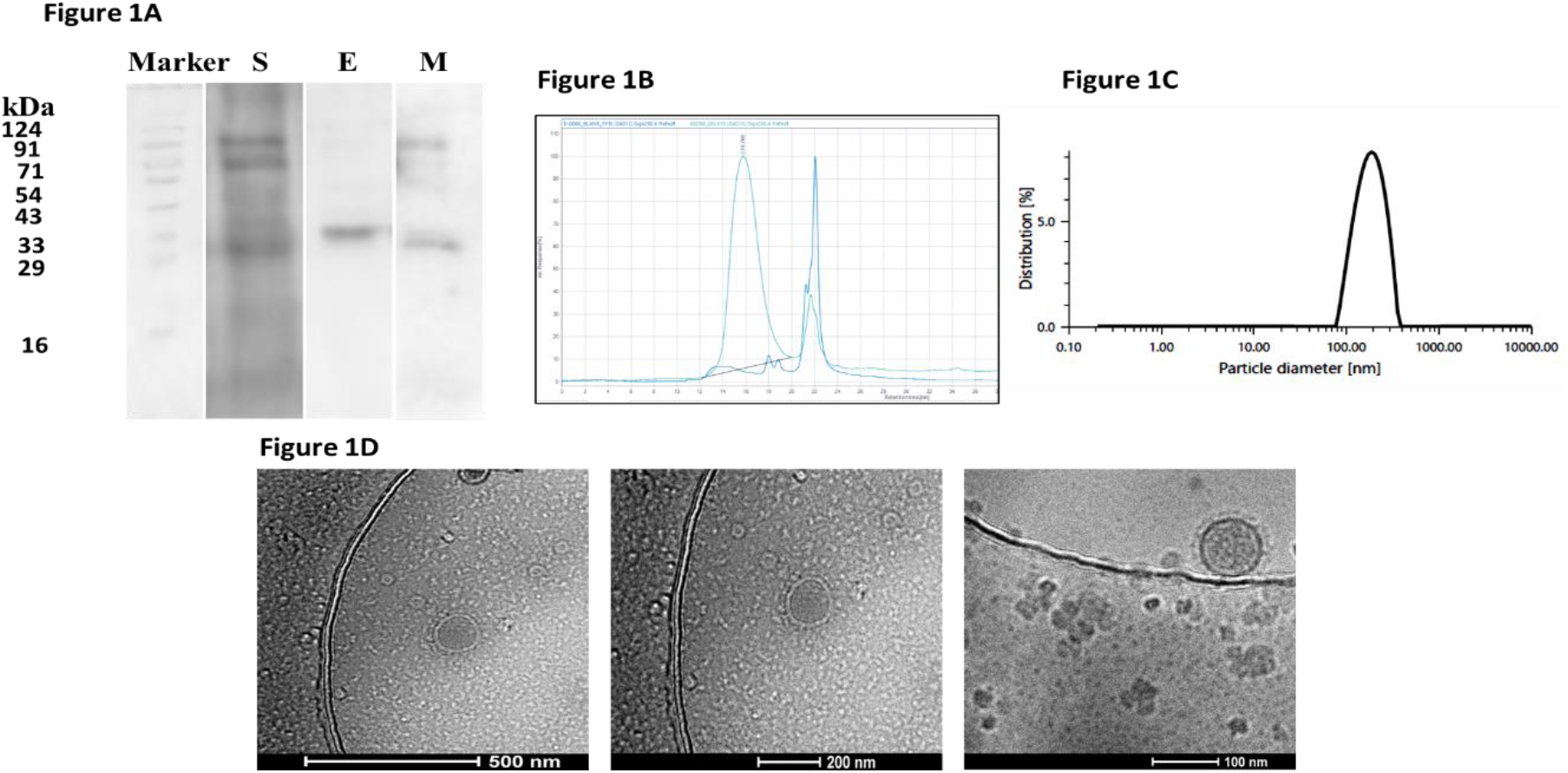
Characterization of the Triple-antigenic PRAK-03202 as a vaccine candidate against SARS-CoV-2 A) Immunoblot analysis for co-expression of Spike, Envelope and Membrane proteins in *S. cerevisiae* based D-Crypt™ platform by using antigen specific antibodies (Sigma Aldrich, USA). M: Marker, S: Spike protein, E: Envelop protein, M: Membrane protein, B) HPLC profile of purified peak (left) from cytoplasmic extract of *S. cerevisiae* cells, co-expressing S, E and M protein at 48 hours post induction. Buffer peak on right side was overlayed with the HPLC profile peak, C) Dynamic Light Scattering (DLS) analysis of the purified VLP Intercept g1^2^ =0.8116, Polydispersity index=18.5%, Particle size=176 ± 18 nm, Baseline=0.998, D) Cryo-TEM image of PRAK-03202 from yeast cell cytoplasmic lysate. The figure shows intact spherical-shaped bilayer particles with distinct spikes.

Binding of the receptor-binding domain (RBD) to ACE2 receptor is the first step in SARS CoV-2 infection. Therefore, preventing the interaction between Spike RBD and ACE2 receptor is considered a strong therapeutic strategy, and work with prior coronaviruses SARS and MERS have demonstrated proof-of-concept for this approach^11,12^. To assess the binding affinity of PRAK-03202 to human ACE2 receptor, flow cytometric analysis was done with Hep-G2 cells (high endogenous expression of ACE2 receptor) and MCF-7 (low endogenous expression of ACE2 receptor-negative control). The data demonstrate that Hep-G2 cells bound preferentially higher (31±9%) than MCF-7 cells demonstrating binding of the PRAK-03202 with the Hep-G2 cells (Figure 2A-B).^13,14^.

**Figure 2.**
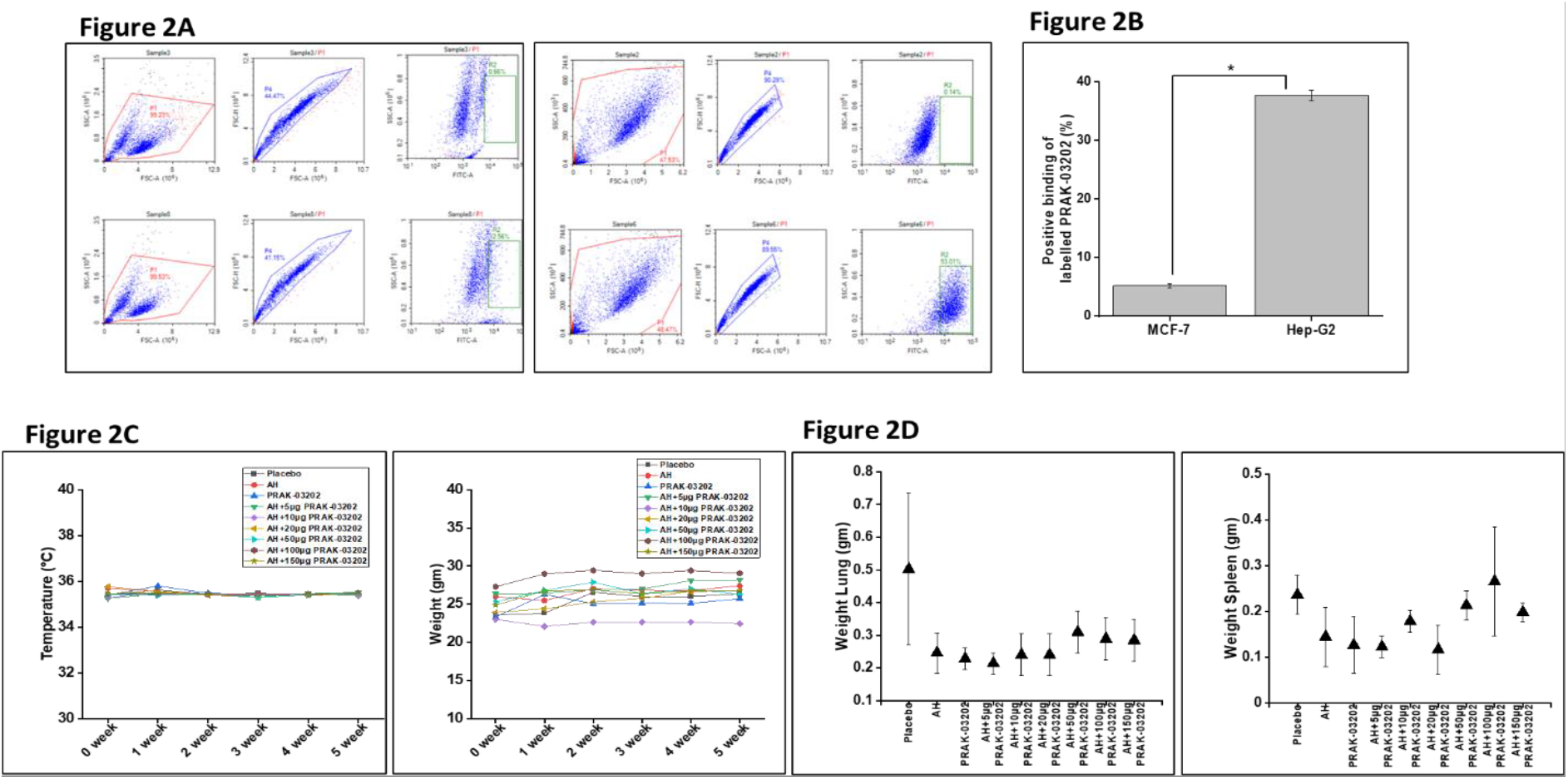
Safety and specificity of PRAK-03202 for ACE2 receptor A) Flow cytometric analysis to show PRAK-03202 binding to the ACE2 receptor on MCF-7 (left panel) and HEP-G2 cells (right panel). PRAK-03202 was labelled with CFSE dye (lower panel); unlabeled PRAK-03202 was taken as control (upper panel). Hep-G2 cells having high endogenous expression of ACE-2 and MCF-7 cells having marginal expression of ACE-2 were used as positive and negative controls, respectively. B) Graphical representation to show positive binding of labelled PRAK-03202 to Hep-G2 and MCF-7 cells. Percentage values are means ± S.E.M. from n = 3. The amount of the binding of the VLP ranged 31±9%. *indicates statistically significant difference (P < 0.05) from MCF-7 cells. Datasets representing many safety related parameters collected during and after immunization such as body weight, body temperature, and organ weight C) Temperature and weight measurement of BALB/c mice immunized with either PRAK-03202 or 6 different doses of PRAK-03202 and alhydrogel (AH) for consecutive five weeks (N=5/group; total number of mice=45) D) Scatter graph depicting weight of lung and spleen of immunized mice at week 5. Values are means± S.E.M.

Serum neutralizing antibodies provide protection against several respiratory viruses and therefore accepted as a functional biomarker of the in vivo humoral response. Therefore, to assess the immunogenicity of PRAK-03202, BALB/c mice (n = 45) were immunized with various doses (5, 10, 20, 50, 100, and 150μg) of PRAK-03202 and categorized into nine groups: Placebo (0μg in physiological saline; N=5), 10μg PRAK-03202 (N=5), alhydrogel(AH) only (N=5), and AH with PRAK-03202 (N=5/ dose; 5,10,20,50,100, and 150μg dose). No visible change in organ/body weight, body temperature (Figure 2 C), or other clinical symptoms, such as an arched back and decreased response of external stimuli even at high doses (150μg) was observed. This dose was, therefore, considered as no observed adverse effects levels (NO-AEL)^15^.

The optimal protection against SARS-CoV-2 is likely to be elicited by humoral and cell-mediated immune responses^16^. Since lower doses in mice are considered to be equivalent to HED^17,18^, next, the total IgG response was evaluated in mice immunized with 5,10, and 20μg dose of PRAK-03202 at days 0 and 35. The total IgG binding endpoint titers (EPTs) from all the immunized mice groups were measured against PRAK-03202 by enzyme-linked immunosorbent assays (ELISAs). This study showed that all the PRAK-03202 vaccinated groups showed a maximum IgG-mediated response than controls. We observed dose-dependent increase in titer throughout the study, with maximum titer obtained by 20μg dose of PRAK-03202 (**AH**: Titer value 1000±0; PRAK-03202: 23500±42; **AH+5μg** PRAK-03202: 8500±50; **AH+10μg** PRAK-03202: 14900±36; **AH+20 μg** PRAK-03202: 25300±65). (Figure 3A). Neutralizing antibody titers (Nabs) of PRAK-03202 closely matches with NAbs titers measured by the convalescent patient sera (51,714±133), highlighting the potential of PRAK-03202 to induce a strong and potent neutralising immune response (Figure 3B).

**Figure 3:**
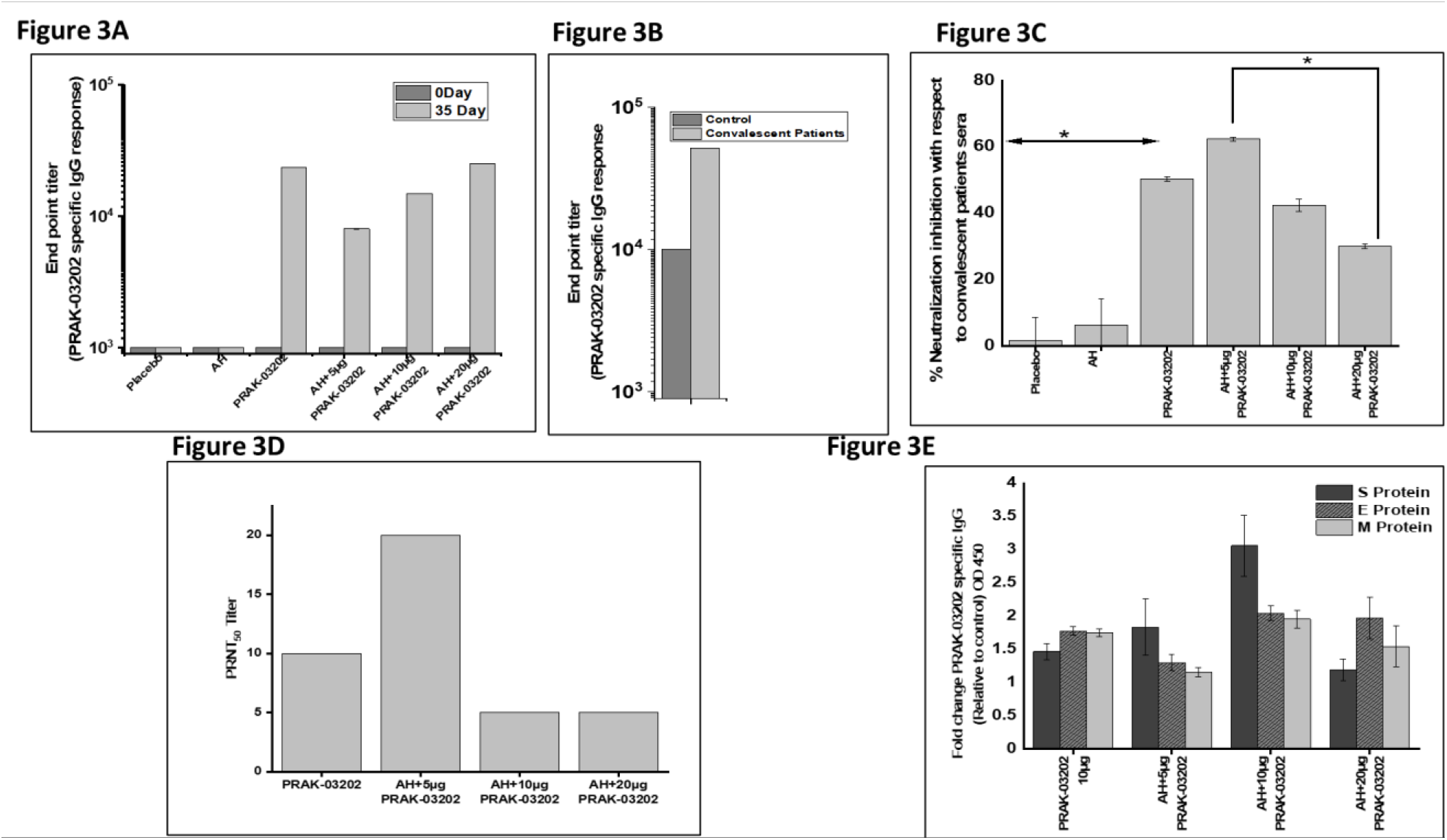
Humoral immune responses in immunized BALB/c mice (n=5/group) and convalescent patients’ sera (N=3). Six groups of BALB/c mice were immunized with indicated doses of PRAK-03202 either with or without AH (n = 5/group). PRAK-03202 specific IgG antibody titers were analyzed by ELISA using HRP conjugated anti mouse secondary antibody (A4416, Sigma, USA) in mice (A) and in (B) healthy and convalescent patients’ sera. Bars are the mean titer with standard deviation indicated. C) Bar chart depicting neutralisation potential (NP) of PRAK-03202 in immunized mice relative to neutralization potential of convalescent patients’ which was taken as reference. Corrected neutralization potential (%) = (Average NP of sera samples of immunized mice/Average NP of sera samples of convalescent patients’) X100. *indicates statistically significant difference (P < 0.05). D) Vero E6 cells were infected with the 30PFU of wild type SARS-CoV-2 virus and serum mixture at the indicated dilutions. PRNT50 is defined as the reciprocal of the highest test serum dilution for which the virus infectivity is reduced by 50% when compared with the average plaque count of the challenge virus control. PRNT50 titer was plotted for each group of immunized mice E) Antigen specific fold change in IgG responses (S, E and M proteins) of PRAK-03202 as measured by ELISA in immunized mice. PRAK-03202 was incubated with 0.25μg of purified S protein, M protein, N protein coated 96 well plate. The plate was read at 450/630nm by ELISA plate reader. Error bars indicate SEM.

RBD-specific neutralization inhibition was evaluated in convalescent patients’ sera and immunized mice. Neutralisation potential (NP) of convalescent patients’ sera was used as a reference and counted as 100 percent; data from mice was plotted relative to that. In the neutralization inhibition assay, sera from all the PRAK-03202 vaccinated groups showed significant NP by inhibiting the RBD and ACE2 interaction compared to control groups. However, maximum inhibition of RBD and ACE2 cell surface receptors was observed at the lowest (5μg) dose of adjuvanted PRAK-03202 (62.3±0.50%) as compared to control groups (AH: 6.12±8.06%; Figure 3C). Comparative low values of total IgG and high values of NP at 5μg dose (NP:62.3±0.50%) suggests that immunization with 10 (NP:42.3±1.92%) and 20μg (NP:29.9±0.69%) of PRAK-03202 and AH leads to generation of more non-neutralizing epitopes in sera that do not engage in ACE2 receptor binding and therefore, less NP was observed. Previous studies have also shown that non-neutralizing epitopes severely reduce vaccine efficacy by inducing strong non-neutralizing immune responses and distracting the host immune system from reacting to the neutralizing epitopes on the RBM^19,20^. Consistent with the previous studies, this study showed that vaccination with lower dose of PRAK-03202 might be more effective against SARS-CoV-2. Recently apparent lack of correlation between IgG ELISA titers with those of the neutralization assay in patients with COVID-1921-24 was discovered. These results suggest that ELISA titers are not a true reflection of NP. We did not find any significant difference in between adjuvanted and unadjuvanted PRAK-03202 (50.3±0.70%) immunized groups. These results demonstrate the binding of the PRAK-03202 induced antibodies to the SARS-CoV-2 RBD, inhibiting the S protein RBD binding to the ACE2 receptor.

Conventionally, neutralizing antibodies are measured by plaque reduction neutralization test (PRNT). PRNT is considered as the gold-standard for determining immune protection^25,26^. In addition, to validate the virus neutralization results, we performed the conventional PRNT on all the PRAK-03202 vaccinated groups. At day 3 post-challenge, wild-type virus–neutralizing activity capable of reducing SARS-CoV-2 infectivity by 50% or more (PRNT50) was detected in all PRAK-03202 vaccinated groups. The serum dilutions yielding 50% virus neutralization (PRNT50 titers) were higher in PRAK-03202 (1:10) and AH plus 5μg PRAK-03202 (1:20) groups in comparison to other adjuvanted groups, where PRNT50 was observed at a dilution of 1:5. Moderate inhibition of plaque number observed in all the mice except mice vaccinated with a lower dose of (5μg) PRAK-03202, suggesting the protective potential of this dose. A strong correlation was observed between the virus neutralization assay and PRNT results, with a correlation efficiency R2 of 0.78(p=0.003) (Figure 3D). The results demonstrate that when diagnosing patient specimens, the viral neutralization assay of PRAK-03202 should deliver results comparable to the PRNT assay, which will require biosafety level 2 (BSL-2) containment instead of BSL3, decrease the turnaround time, and increase the testing efficacy to high throughput in case of SARS-CoV-2 infection.

To further investigate the immunogenicity of PRAK-03202, detailed immune responses induced by PRAK-03202, including SEM specific immune responses, were observed, which have not been investigated previously. Spike, envelope, and M-specific (SEM) antibody responses were evaluated by ELISAs at week five after the initial immunization. IgG response is observed against all three antigens in animals vaccinated with PRAK-03202 when compared with controls (Figure 3C), which suggest immunogenicity towards all the three antigens (Spike IgG-1.46±0.11 fold, Envelope IgG-1.77±0.06fold, Membrane IgG-1.74±0.056 fold) (Figure 3E).

Evaluation of immune response against the S, E and M showed that mice vaccinated with 10μg dose showed maximum immune response against the three antigens (S-3.05±0.45 fold, E-2.04±0.11-fold, M-1.95±0.13 fold; P=NS) while alhydrogel in combination with 5μg PRAK-03202 showed comparative high response for spike protein when compared with controls (AH and 5μg PRAK-03202: S-1.83±0.42fold, E-1.35±0.12 fold, M-1.3±0.07fold). (Figure 3E). This highlights the potential of PRAK-03202 to induce a strong and potent immune response by eliciting effective antibody responses for all three antigens and it could serve as an effective vaccine candidate for novel coronavirus infections.

To further observe that PRAK-03202 is being recognised by the sera of convalescent patients(n=7), Western blot analysis was performed. Our-findings showed multiple protein bands, which speculate that certain components of PRAK-03202 are being recognised by antibodies present in sera of convalescent patients’ (Figure 4A).

**Figure 4.**
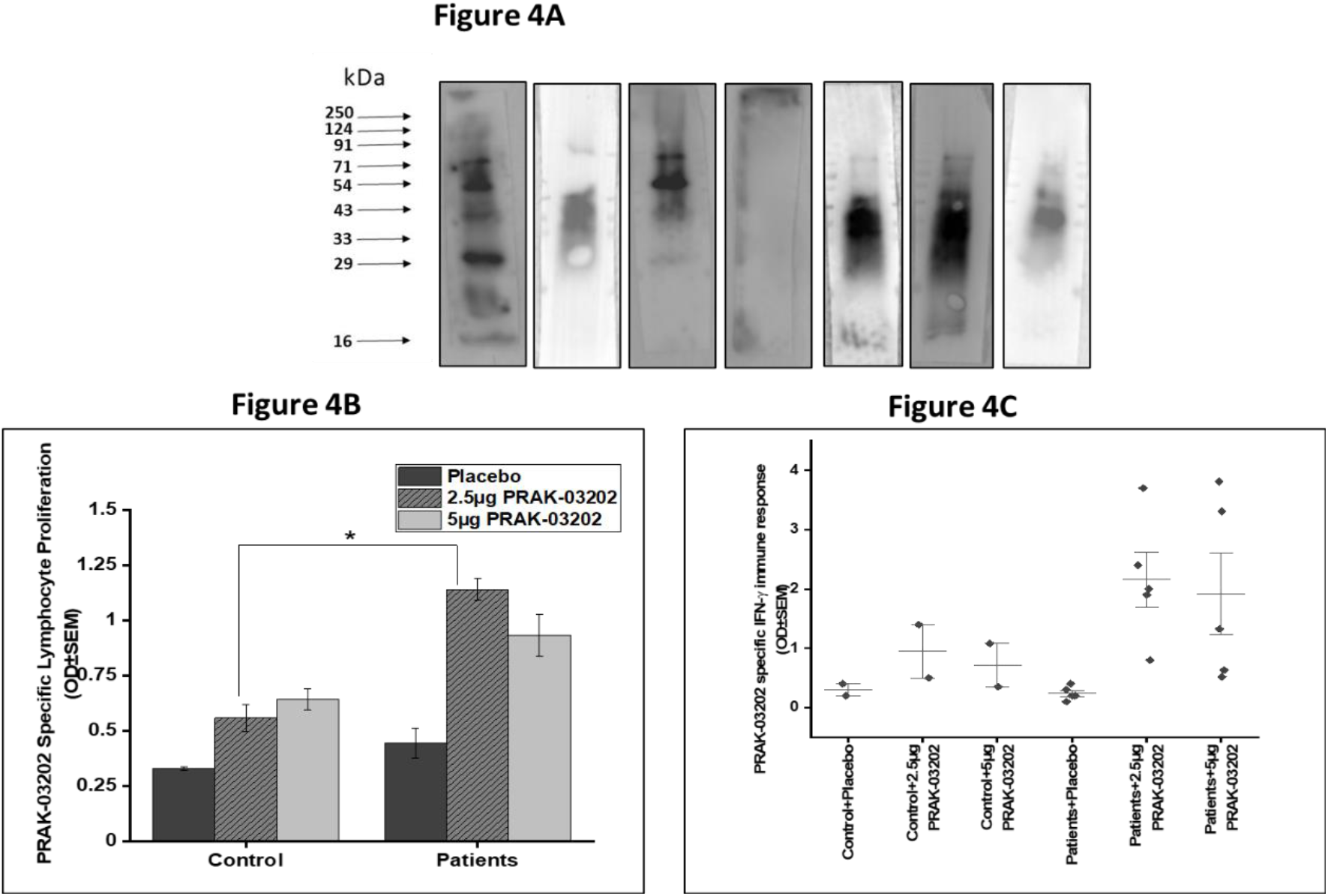
Cell mediated immune responses in convalescent patients A) Immunoblot analysis of PRAK-03202 when probed with serum of convalescent patients’ sera (n=7). M=marker, P1-P7: Convalescent patients’ samples B) Lymphocyte proliferation in PRAK-03202 (2.5μg and 5μg) stimulated PBMCs of controls (n=2) and convalescent COVID-19 patients (n=5). Samples were stimulated for 120 hours and BrDU assay was performed (Cell Proliferation ELISA BrDU, Roche, USA). Plate was read at 450/690nm by ELISA plate reader. OD values are means ± S.E.M. Significant difference in between controls and patients was observed. *indicates statistically significant difference (P < 0.05) C) Cytokine staining for IFN-γ by ELISA (Human IFN-γ ELISA set (RUO), BD Biosciences, USA) in controls and patients. Both 2.5μg and 5μg dose of PRAK-03202 showed elevated levels of IFN-γ in patients in comparison to controls. Error bars indicate SEM.

Not just humoral responses, cellular immune responses have also been shown to be associated with more favourable recovery in MERS-CoV infection and are likely to be important against SARS-CoV infection27. To assess T cell responses and if the PRAK-03202 mimics the viral epitopes, PRAK-03202 was used in cytokine staining assays from healthy and convalescent human PBMCs. PRAK-03202 specific lymphocyte proliferation was evaluated by ELISA in convalescent patients (N=5) and healthy controls sera (N=2).

In comparison to controls, higher lymphocyte proliferation was observed in PRAK-03202 stimulated patient’s sera at two doses (2.5μg-2 Fold; 5μg:1.45 Fold) (Figure 4B). The presence of PRAK-03202 thus suggests that it induces lymphocyte proliferation in convalescent blood, indicating its recognition by APCs and presentation to immune cells.

The dominance of type 1 or type II [4-7] immunity is elicited by vaccines in previous studies. Therefore, we evaluated the IFN-γ response in these patients (Figure 4C). Experiment data showed elevated IFN-γ levels in PRAK-03202 stimulated patients PBMCs (2.16±0.33) as compared to healthy controls (0.95±0.23), suggesting induction of T helper 1 (Th1)–biased cellular immune responses. Rapid cellular responses could potentially reduce the viral load & spread of SARS-CoV-2 and the associated COVID-19 illness. These findings support the previous studies demonstrating intranasal IFN-γ as a viable treatment option for Respiratory syncytial virus (RSV) due to its capability of reducing viral titers in the lung with no detectable increase in CD4+ or CD8+ T cell infiltration^28^. This indicates the potential mimicking of PRAK-03202 to the actual virus.

Safety and efficacy are essential for vaccine development at both preclinical studies and clinical trials. T-cell infiltration could increase lung viral titers in the case of respiratory viruses and is considered a major culprit in vaccine development for respiratory diseases^28^. We provide evidence for the safety of PRAK-03202 in mice and did not observe any visible change in lymphocyte subset percentage (CD4+, CD8+, and CD11b+, NK, Ly6B cells) in either lung (Figure 5) or blood (Figure 6A) of PRAK-03202 vaccinated groups. This speculates that PRAK-03202 could be an effective treatment option for SARS-CoV-2.

**Figure 5.**
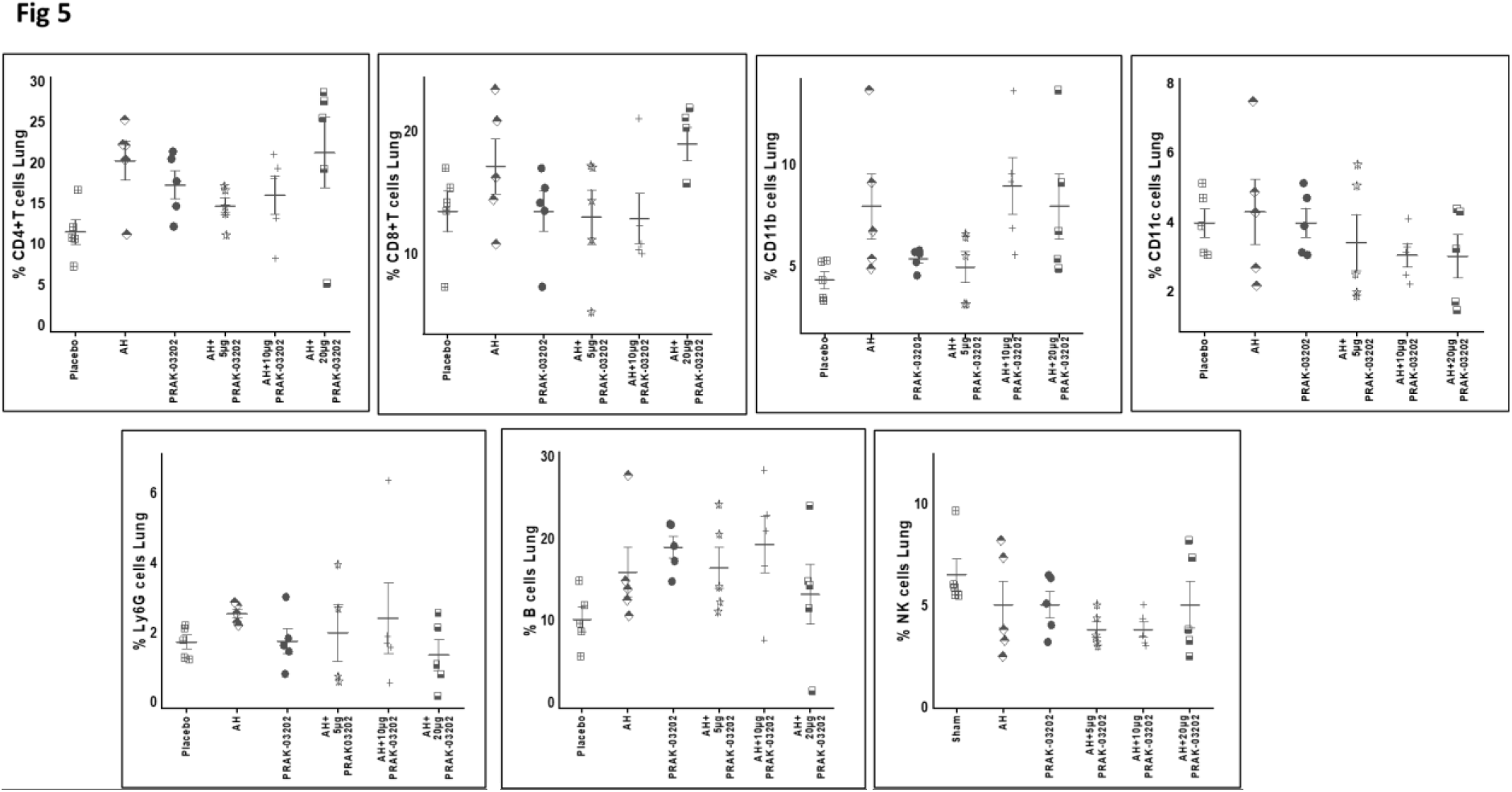
Profile of immune cells in lung of vaccinated BALB/c mice with PRAK-03202. Lungs were initially treated with collagenase solution and macerated. Flow cytometric analysis of lymphocyte subset percentage was performed by using specific antibodies for indicated antibodies at 1:100 dilution. T-cell infiltration could increase lung viral titers in case of respiratory viruses and is considered major culprit in treating respiratory diseases. No visible specific stimulation and marginal infiltration of T-cells suggest vaccine efficacy of PRAK-03202. Error bars indicate means ± S.E.M.

**Figure 6.**
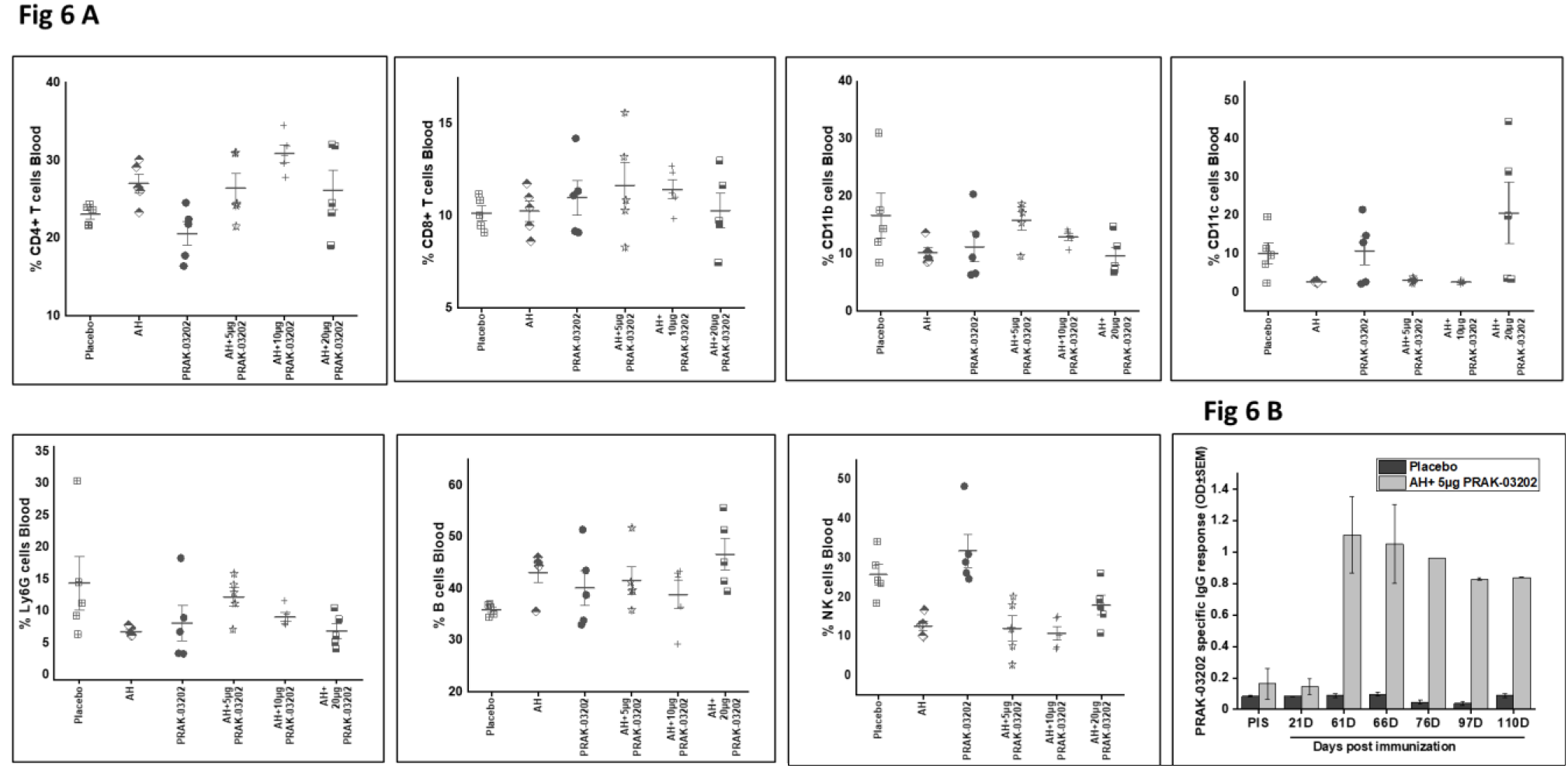
Profile of immune cells in blood of mice vaccinated with PRAK-03202. A) Flow cytometric analysis of lymphocyte subset percentage was performed by using antibodies for the indicated molecules at 1:100 dilution. No visible specific stimulation of blood cells and marginal infiltration of T-cells suggest that PRAK-03202 could be a better therapeutic option for SARS-CoV-2. Error bars indicate means ± S.E.M. B) Sera from immunized mice were collected at various time points (0,21,61, 76, 86, 97 and 110 days). Total IgG response was evaluated by enzyme-linked immunosorbent assays (ELISAs) using 1:5000 dilution of HRP conjugated secondary antibody (A4416, Sigma, USA). IgG response even after 110 days suggest long term efficacy of PRAK-03202. Error bars indicate SEM

Mice vaccinated with adjuvanted PRAK-032025 were kept alive beyond the recovery period under observation for long term humoral immune response; sera were kept at various time points. Total IgG responses could be seen even after 110 days (IgG: 5.25±0.23 Fold as compared to placebo) suggesting, long-term protective efficacy of PRAK-03202 (Figure 6B).

The novel coronavirus, SARS-CoV-2, which has affected at least 114 countries and killed more than 864,000 people, has been officially declared a pandemic by WHO. The incidence of mortality and morbidity observed with this pandemic has created an imminent need for a prophylactic vaccine to prevent ~7 Billion people worldwide. Though several vaccine candidates have initiated clinical testing, very little is currently known about their protective efficacy^29,30^. Here preclinical development of a VLP vaccine candidate, PRAK-03202, to combat this emerging infectious disease is described. To produce PRAK-03202, we have used the D-crypt^TM^ platform due to its low risk of contamination by adventitious agents, low production costs, and the ability to produce VLPs with reliable qualities^9^.

This study further demonstrates the neutralization potential and antigen-specific immunogenicity, and cell mediated immunity of the PRAK-03202 vaccine candidate. A strong correlation between the neutralization and PRNT50 assay could fill a major gap for COVID-19 surveillance and vaccine development by requiring BSL-2 containment, decreasing the assay turnaround time, and increasing the testing efficacy to high throughput scale.

PRAK-03202 provides several advantages over other vaccine platforms including 1) resemblance with native SARS CoV-2, 2) Non-replicative and non-infective nature, making them safe to use in immunocompromised populations^31^, 3) Scalability and commercialization using our D-crypt^TM^ platform will circumvent conventional vaccine production complexities, and thus represent an attractive vaccine strategy against SARS-CoV-2 infection. Previous studies on SARS CoV VLPs were focused on the assembly of virions and did not investigate their immunogenicities ^32^. We report here on the development of PRAK-03202, its immunogenicities, and its potential as an anti-SARS vaccine. Our results showed that PRAK-03202 could induce a cellular immune response, neutralizing antibodies against SARS CoV-2, and is safe in nature. Additionally, neutralizing PRNT50 average responses in live virus challenge assay speculate protection against the SARS CoV-2 challenge with PRAK-03202. These results collectively suggest making to the clinical development of PRAK-0302 for use in humans. These results support the further evaluation of PRAK-03202 as a vaccine candidate for COVID-19 in clinical trials.

## Methods

### Cloning and expression of “S,”, “E,” and “M” proteins

The three proteins S, E, and M were co-expressed in protease deficient D-Crypt^TM^ platform using vectors, pYRE100 and pYRI10033. The M protein was cloned in pYRI100 integration vector (pYRI100_CMP), while S and E (pYRE100_CSP_CEP) protein were cloned into pYRE100 episomal single expression vector as two separate expression cassettes for co-expression analysis. The vector pYRE100 plasmid was also transformed as control. Transformation was performed by using Lithium acetate/SS-DNA/PEG mediated protocol. The synthetic genes sequence biased and optimised for expression in *S. cerevisiae* host was obtained from GeneArt (Regensburg, Germany).

### Transformation in D-Crypt^TM^

For co-expression of three proteins, the M protein gene (pYRI100_CMP) was integrated into protease deficient proprietary *S. cerevisiae* host genome (PYPD), and transformants were selected over selective YNB Glucose without LEU auxotrophic marker plates. This was followed by the transformation of the S and E protein genes (pYRE100_CSP_CEP episomal construct) in PYPD strain integrated with the M gene construct. Transformants were selected over selective YNB Glucose without URA and LEU auxotrophic marker plates. Vector pYRE100 plasmid was transformed in protease deficient *S. cerevisiae host* as control and was selected over selective YNB Glucose without URA auxotrophic marker plates. Sequential transformation was performed using Lithium acetate/SS-DNA/PEG mediated protocol by incubating the plates at 28°C for 2-4 days for S, E, and M protein co-expression analysis.

### Expression analysis of culture

Cultures were grown for the co expression of three proteins in YNB Glucose without URA and LEU media at 28°C for 36 hr and control with URA media. At this stage, cultures were harvested and induced with galactose at a final concentration of 2% in YNB minimal medium for 48 hours. Following induction, the harvested culture was used for expression analysis.

### Immunoblot

In order to confirm the presence of the S, E, and M protein, immunoblot analysis was carried out using the purified PRAK-03202. For immunoblot, ~6 μg samples were loaded on SDS-PAGE under reduced conditions. Immunoblot was developed using 1:1000 dilution of S specific antibody (Cat≠ZHU1076, Sigma-Aldrich) and 1:500 dilution of polyclonal M (Cat≠ GTX134866, Gene Tex) and E (Cat≠MBS150849, My Bio-source) specific antibodies. Signals were detected with ECL solution (Biorad cat#170-5061) using the Western blot development instrument (G: Box, SYNGENE, UK).

### SEC-HPLC analysis

100μL of PRAK-03202 sample was injected onto Agilent Bio SEC-5 (5μ, 2000Å, 7.8 mm ID × 300mm) (Agilent Technologies) column for analytical SEC analysis and analysed with Agilent Technologies HPLC instrument (1260 Infinity II). The column is packed with 5μm silica particles coated with neutral hydrophilic layer for maximum efficiency and for better separation. The estimated exclusion limit of this column for proteins was 10,000,000 Da for globular proteins. 20 mM sodium phosphate, pH 7.0 with 400 mM sodium chloride, was used as a standard SEC buffer for the elution, which provides necessary ionic strength to the PRAK-03202. UV signals were traced at 214 nm and 280 nm. 0.5 mL/min was the flow rate used for the analysis.

### Dynamic Light Scattering Analysis

Dynamic light scattering analysis was done to analyse the size and homogeneity of purified PRAK-03202. Measurements were taken on a Litesizer 500 (Anton Paar, Austria) instrument in automatic mode. The size of the molecule was measured as the function of maximum intensity observed. The laser used to measure the hydrodynamic diameter had a wavelength of 658nm. During the measurement, an automatic mode was selected for the light scattering angle, which automatically selects the angle based on the particle concentration. In this case, side scattering (90°) was used by the instrument to measure the Hydrodynamic diameter.

### Electron microscopy

To characterise and determine the size and structure of PRAK-03202, the HPLC purified fractions from the cell lysate were subjected to electron microscopic analysis, as described previously^9^. The fractions were briefly adsorbed onto Carbon-Formvar copper grids and subsequently negatively stained with 1.5% uranyl acetate aqueous solution for the 50s. Subsequently, the grids were washed and examined on the Talos Arctica transmission electron microscope.

### Binding affinity of PRAK-03202

Flow cytometry analysis was performed to detect the binding of PRAK-03202 to hACE2 receptor in Hep-G2 cells and MCF-7 cells. Previous studies support the higher expression of ACE-2 in Hep-G2 cells^34^, while marginal expression of ACE-2 receptor could be seen in MCF-7 (https://www.proteinatlas.org/ENSG00000130234-ACE2/cell). 3 μg of purified PRAK-03202 was labelled with 5 μM of CFSE (Thermo, C34554) for 3 hrs at room temperature, which was followed by incubation with Hep-G2 and MCF-7 (5 x 10^4^ cells/100 100 μl) cells for 1 hr on ice). After 1 hr of incubation, samples were transferred to 96 well-round plate and washed twice with FACS buffer (PBS+ 2%FBS) by centrifuging plates at 1000 rpm for 5 mins at 4°C. Samples were acquired and analyzed by flow cytometry (Novocyte, ACEA) for binding analysis using the FITC channel.

### Mice

Specific pathogen-free 6-8 weeks-old BALB/c mice were maintained in the DHITI Life Sciences, India. All animal protocols were approved by the Committee for the Purpose of Control and Supervision of Experiments on Animals (1931/PO/RcBi/S/16/CPCSEA). BALB/c mice (n = 45) were immunized with PRAK-03202 at 14 days of interval and categorized into the following nine groups: Placebo (0μg in physiological saline; N=5), 10μg PRAK-03202 (N=5), alhydrogel(AH) only (N=5), and AH with PRAK-03202 (N=5/dose; 5,10,20,50,100, and 150μg per dose).

### Clinical information and serological testing

Patients’ medical history was taken for information on demographic characteristics, the clinical profile including blood biochemistry, lung profile, renal profile, liver profile, and subsequent outcome in case of any reported symptoms and signs. Blood from the convalescent patients was fractionated into sera and PBMCs. The study is approved by the Institutional Ethics committee at Premas Biotech Pvt. Ltd.

### BrDU assay

PBMCs were separated from the blood of the convalescent patients and stimulated with two doses of PRAK-03202 (2.5μg and 5μg). The proliferation rate of lymphocytes was measured using 5-Bromo-2-deoxyuridine (BrDU) assay as per the manufacturer instructions (Cell Proliferation ELISA BrDU, Roche, USA).

### Serum Antibody Measurements

IgG mediated antibody titer of serum samples collected from immunized animals and convalescent patients’ sera were determined by enzyme-linked immunosorbent assay (ELISA)35 using 1:5000 dilution of HRP conjugated anti mouse (A4416, Sigma-Aldrich, USA) and 1:10000 dilution of HRP conjugated anti human secondary antibody (A0170, Sigma-Aldrich, USA) respectively. For antigen-specific IgG response, 96-well microtiter plates were coated with 0.0.25μg of purified S protein, E protein, M protein individually at 2-8°C overnight and blocked with 2% BSA for 1h at room temperature. Diluted sera (1:1000) were applied to each well for 2h at 37°C, followed by incubation with HRP conjugated goat anti-mouse antibodies for 1h at 37°C after 3 times PBS wash. The plate was developed using TMB, following 2M H_2_SO_4_ addition to stop the reaction, and read at 450/570 nm by ELISA plate reader for final data.

### SARS-CoV-2 neutralization assays

Neutralization experiments on serum samples of immunized animals and convalescent patients were performed as per the manufacturer protocol (sVNT Kit, GenScript, USA). Briefly, the samples and controls were pre-incubated with HRP-RBD to bind the circulating neutralizing antibodies to HRP-RBD. The mixture was added to an hACE2 protein pre-coated plate and incubated for 15 minutes at 37°C. After 4 times washing, the mixture was incubated with TMB for 15 minutes at 20-25°C in the dark. The stop solution was added, and the plate was read at 450nm in a microtiter plate reader. Neutralisation potential of mice was plotted relative to human neutralization potential (reference), taken as 100 percent.

### Plaque reduction neutralization test (PRNT)

Vero E6 cells (1×105 per well) were seeded to 24-well plates. On the following day, 30 PFU of infectious wild-type SARS-CoV-2 was incubated with diluted serum (total volume of 150μl) at 37°C for 1 h. The virus-serum mixture was added to the pre-seeded Vero E6 cells and incubated at 37°C for 1 hour. Following this, 1 ml of 2% Modified Eagle’s Medium (MEM) containing carboxy methylcellulose (CMC) was added to the infected cells. After 3 days of incubation, 1 ml 3.7% formaldehyde was added and incubated for 30 minutes at room temperature. The formaldehyde solution from each well was discarded, and the cell monolayer was stained with a crystal violet solution for 60 minutes. After washing with water, plaques were counted for PRNT50 calculation. The PRNT assay was performed at the BSL-3 facility at THSTI, India.

### Cytokine profiling

Healthy controls and patients PBMCs (1×105 cells/well) were stimulated with two doses of PRAK-03202 (2.5μg and 5μg) for 120 hrs at 37°C in a humified chamber containing 8% CO_2_. Supernatants were collected, and cytokine staining for IFN-g was performed as per the manufacturer instructions (Human IFN--g ELISA set (RUO), BD Biosciences, USA).

### Vaccine safety evaluation

SARS-CoV-2 trial vaccine’s safety was evaluated in immunized mice at week 5. Datasets of many safety-related parameters were collected during and after immunization, including clinical observation, body weight, organ weight (lung and spleen), body temperature. Flow cytometric analysis of lymphocyte subset percentage was performed in lung and collected blood by using commercially labelled CD4+ (Cat≠100412; Biolegend):, CD8+ (Cat≠140410; Biolegend), CD11c (Cat≠117316; Biolegend), CD11b+ (Cat≠101224;Biolegend), B cells (Cat≠1115535; Biolegend), NK (Cat≠108905; Biolegend), Ly6B (Cat≠127629; Biolegend) antibodies. Briefly, the lungs were initially treated with collagenase solution and macerated. Both lung and blood cells were lysed in RBC lysis buffer (Biolegend) for 5 minutes on ice. Following lysis, blood cells were centrifuged at 400rpm for 5 minutes at 4°C. Cell pellets were resuspended in FACS buffer containing 1:100 dilution of target-specific antibodies and incubated for 20 minutes at 4°C in the dark. After centrifugation and two times washing with FACS buffer, data were acquired using a BD FACS Canto II and analyzed using FlowJo (TreeStar).

### Long term humoral immune response

Mice have been kept alive beyond the recovery period under observation for immune response and long-term immune response. Briefly, sera were collected from these mice at various time points (0,21,61,66,76,86,97 and 110 days). Total IgG response was evaluated by enzyme-linked immunosorbent assays (ELISAs) using 1:5000 dilution of HRP conjugated anti mouse secondary antibody (A4416, Sigma-Aldrich, USA).

### Statistical Analysis

Results are presented as mean ± standard error of the mean (S.E.M.). The difference in antibody titers was evaluated by the non-parametric Kruskal-Wallis test. The difference in neutralization potential and overall differences were tested using analysis of variance (ANOVA). Bonferroni-adjusted P-values were computed for all pair-wise comparisons. The correlations between methods were analyzed by Spearman correlation test. All the statistical analysis was performed using OriginPro (Version:2020b). A P value less than 0.05 was considered statistically significant.

## Author information

### Contributions

NM, SM, RR and PKK designed and planned the animal study. KA developed D-Crypt™ and strategy for clone development. AU, PKK, AJ and BP contributed to the purification development of PRAK-03202. BP and DK performed all the ELISA experiments. VBA, IS, PF and BK performed the 5L fermentation scaleup studies. SK, SP and SSR performed all the analytical studies (DLS, HPLC, Western blot). SM, RR, AM, AG MKS and KJ performed animal studies. DP, JP SKS AN AS, DK SB, SR contributed to the clone development. RG analyzed the data with input from all authors.

## Acknowledgments

The authors thank Dr Manidipa Banerjee, IIT Delhi for TEM analysis, Dr. Deepak Rathore and Vishal Gupta at THSTI Faridabad for FACS analysis, Dr. Sankar Bhattacharyya, THSTI for PRNT assay, Dr. Nipendra Singh at RCB for mass spectrometry, Rakesh Jaswal and Abhishek Choudhary for logistics. We would also like to thank Dr. Manish Kumar and Nupur Pandey, Premas Biotech, for providing their assistance while performing ELISA.

The work was funded by Akers Biosciences and Premas Biotech Private Limited, India.

