## Supplemental Data for "PRAK-03202: A triple antigen VLP vaccine candidate against SARS CoV-2"

Figure 1A

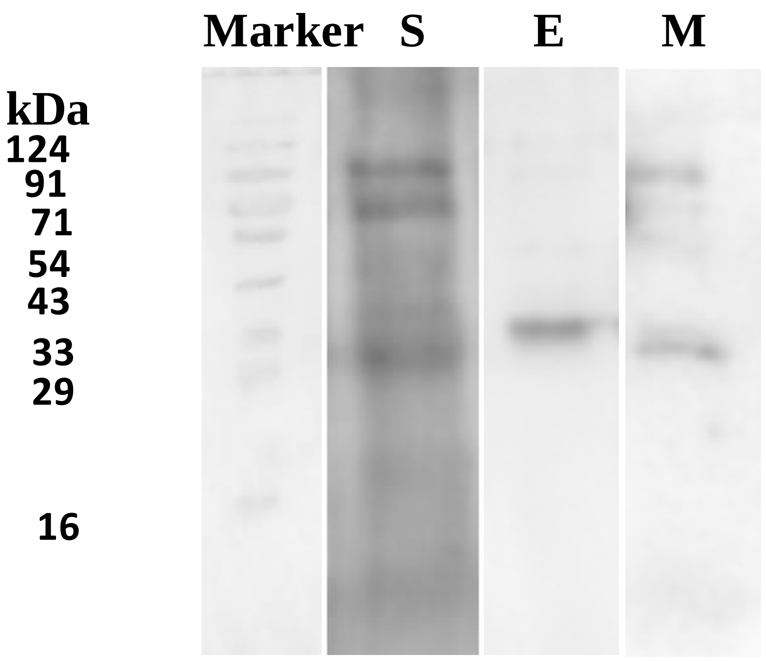


Figure 1B

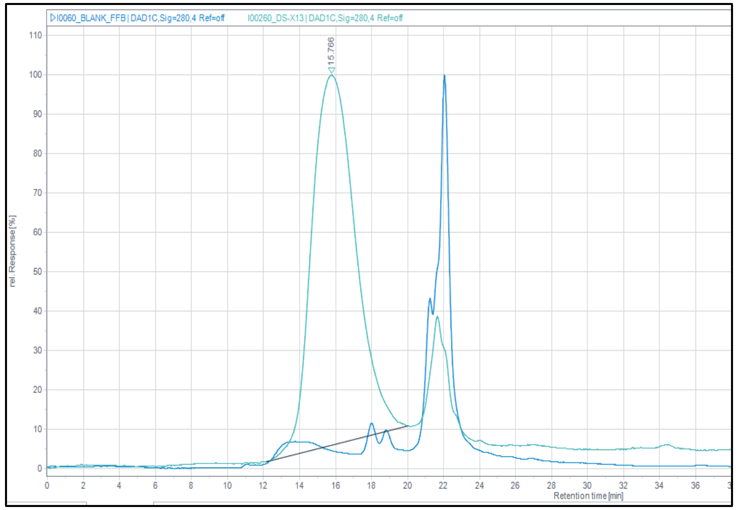

Figure 1C

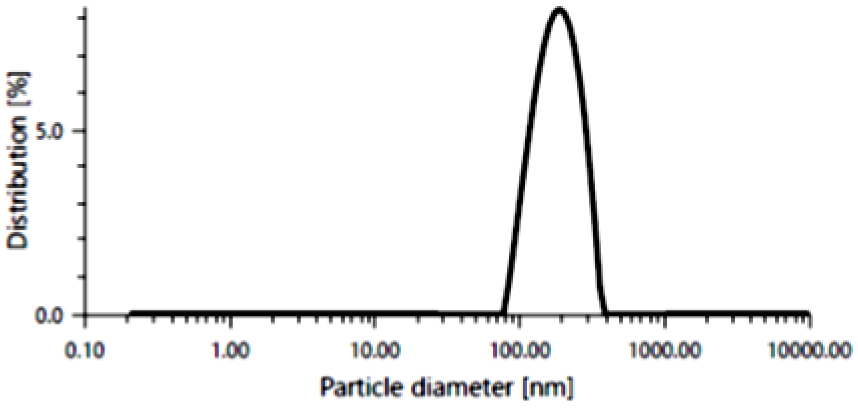


Figure 1D

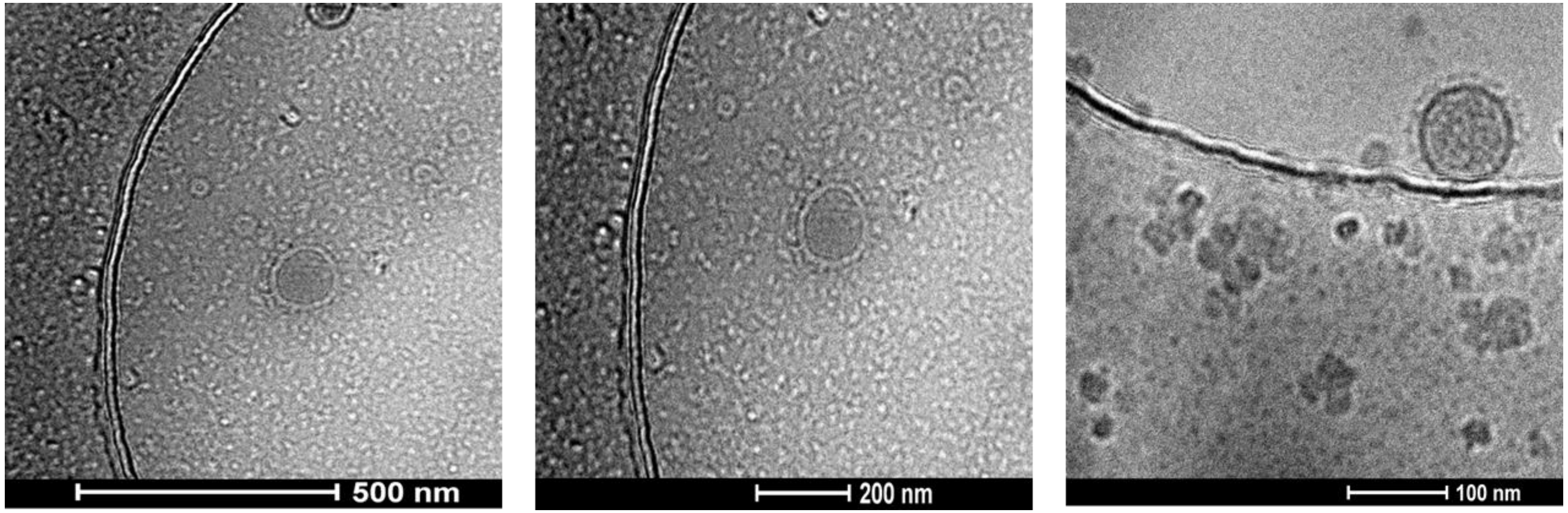

### Figure 2A

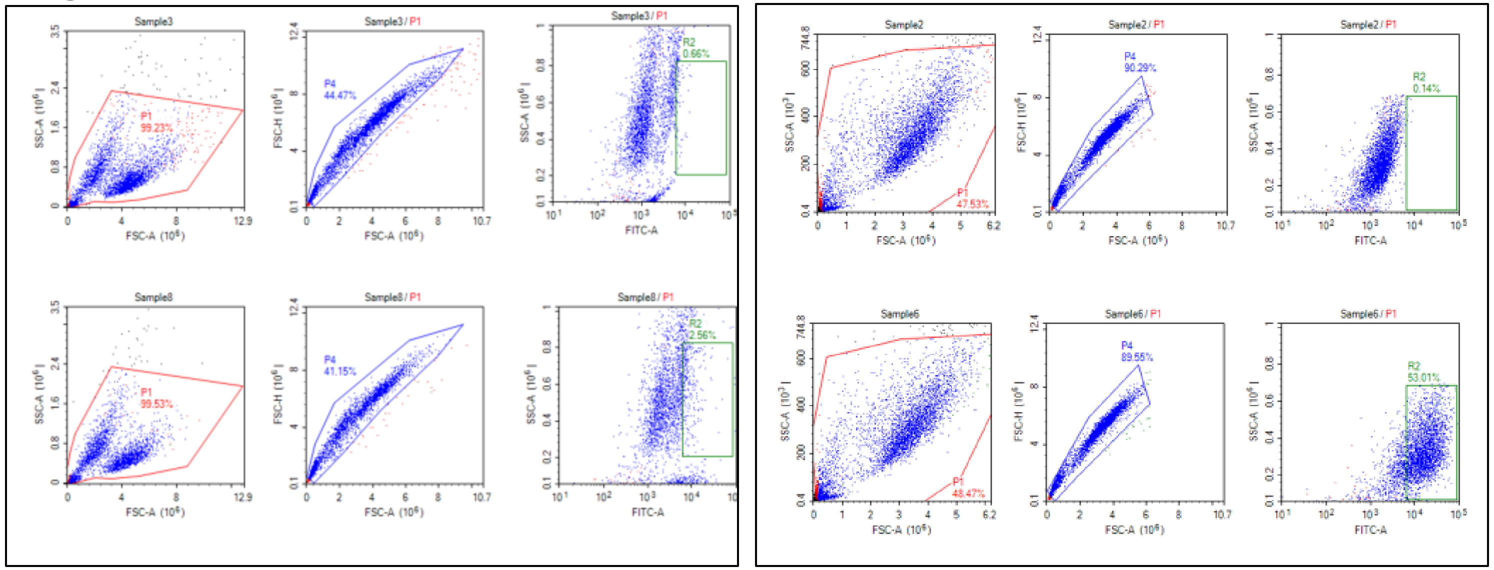


### Figure 2B

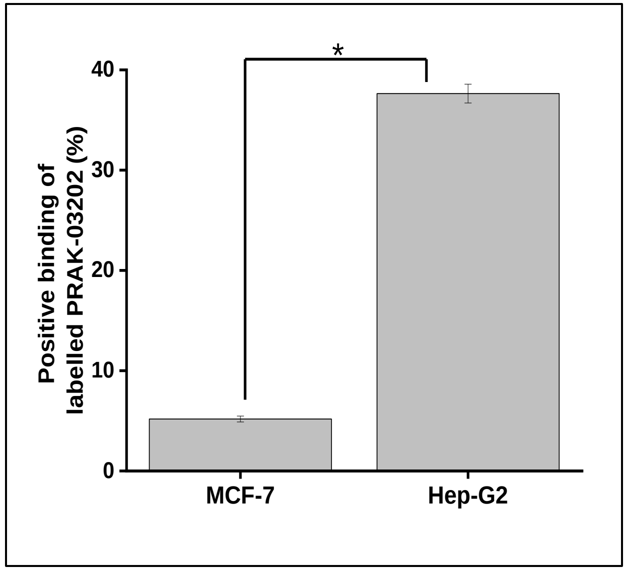


### Figure 2C

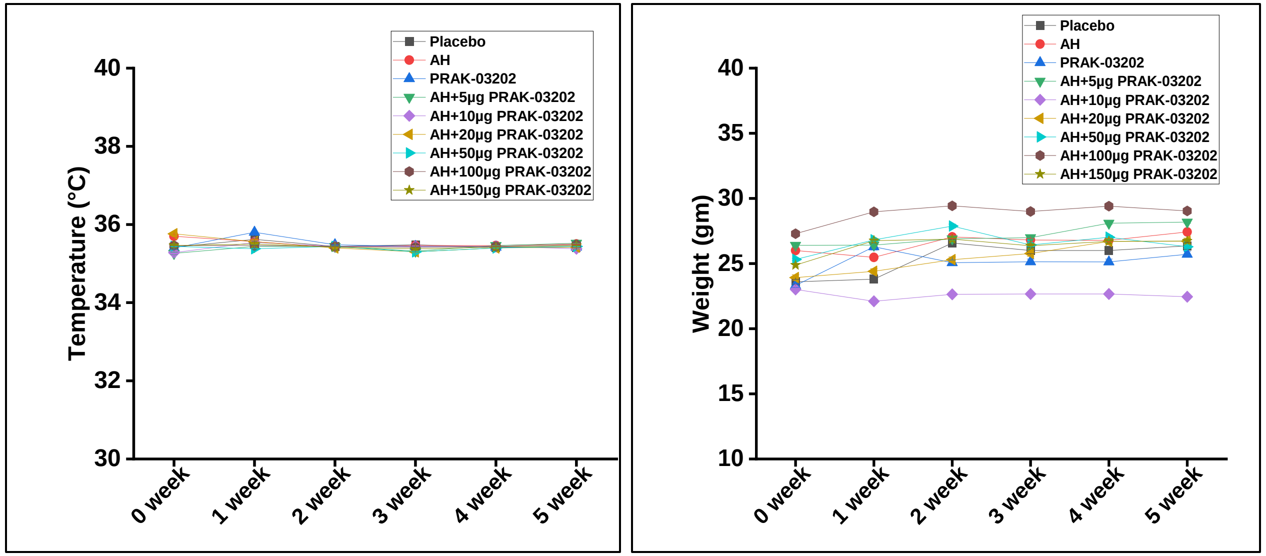


### Figure 2D

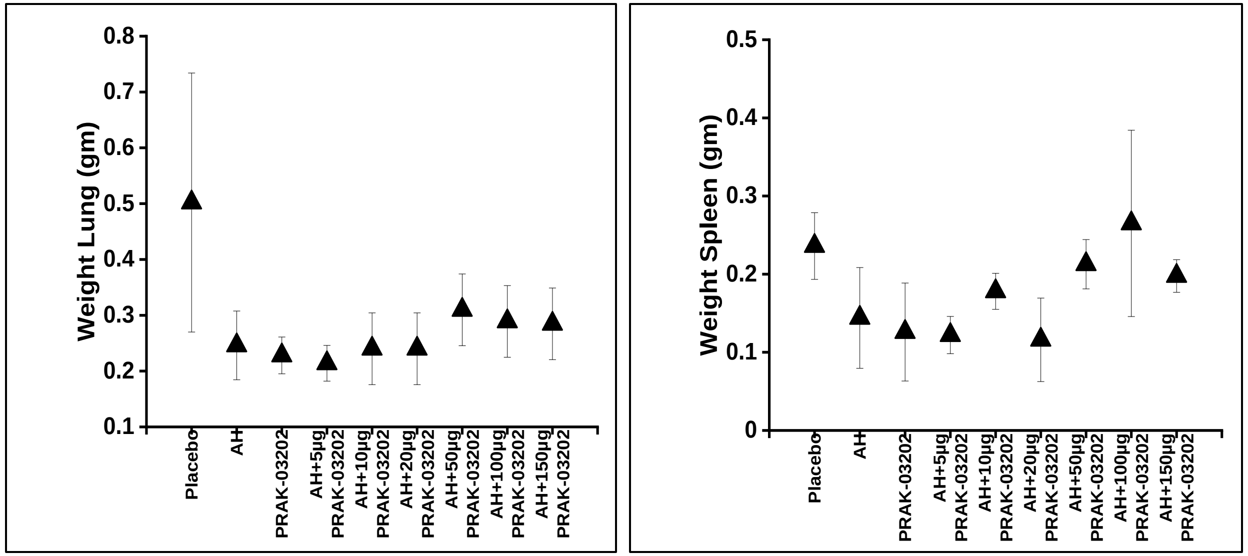

Figure 3A

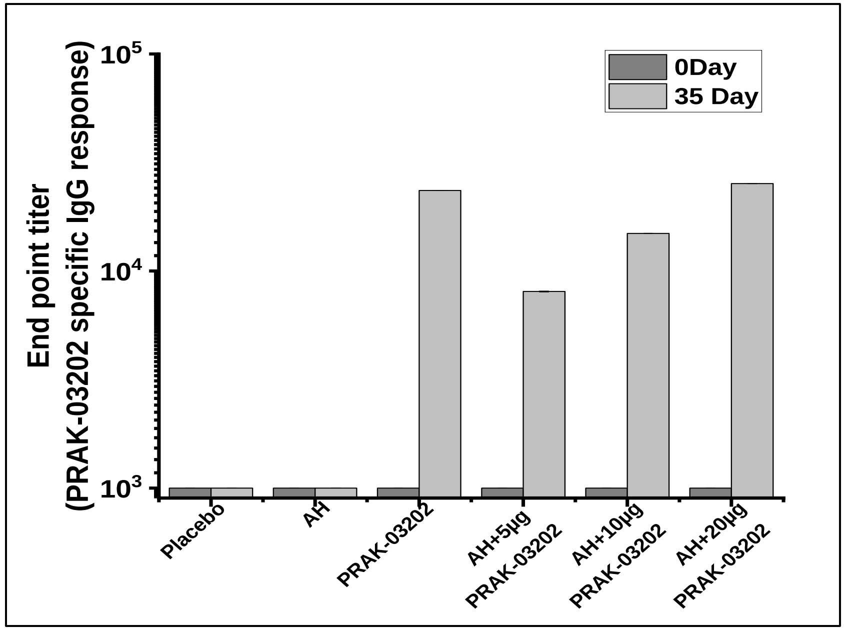


Figure 3B

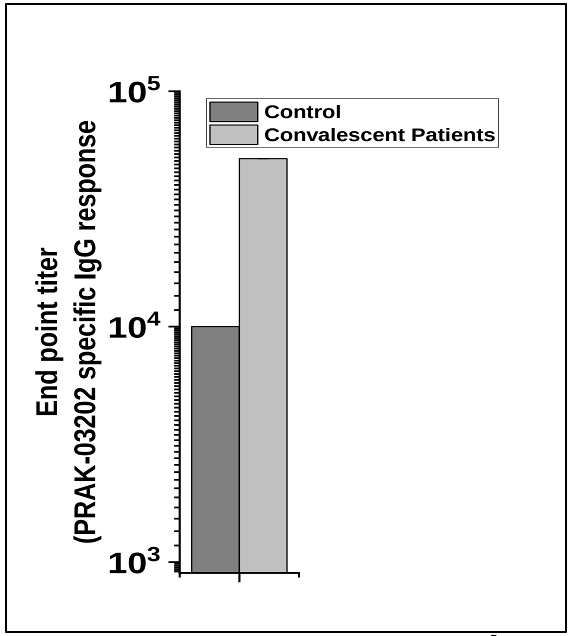


Figure 3C

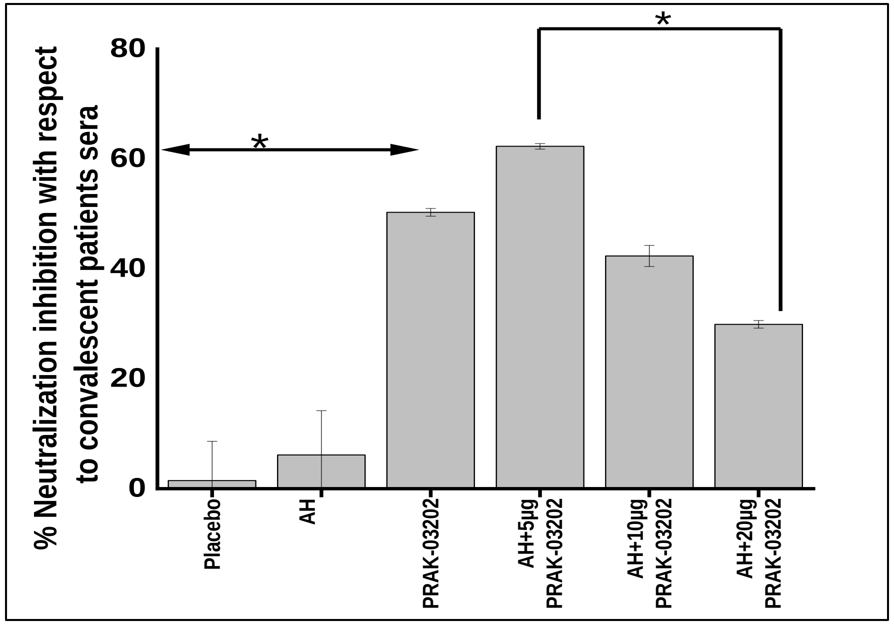


Figure 3D

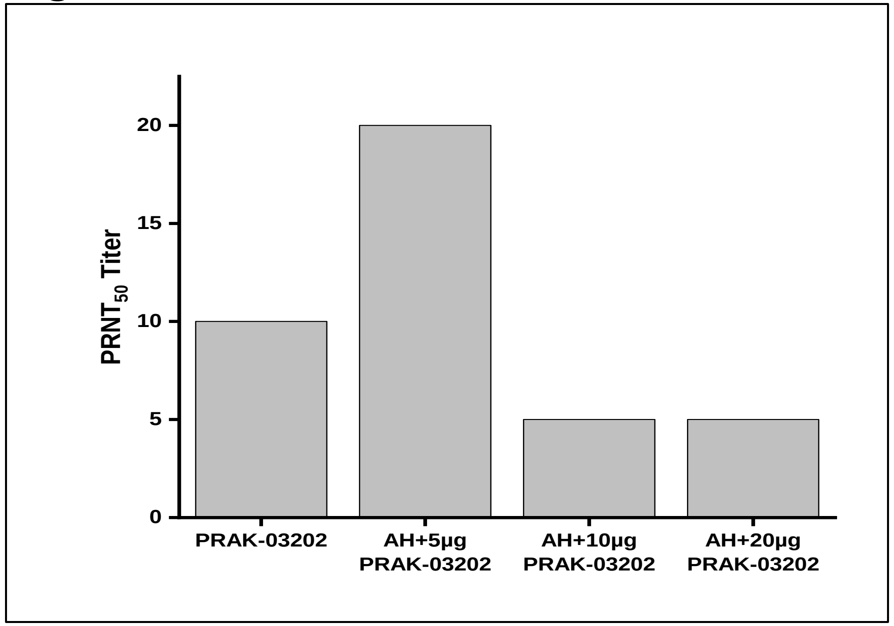


Figure 3E

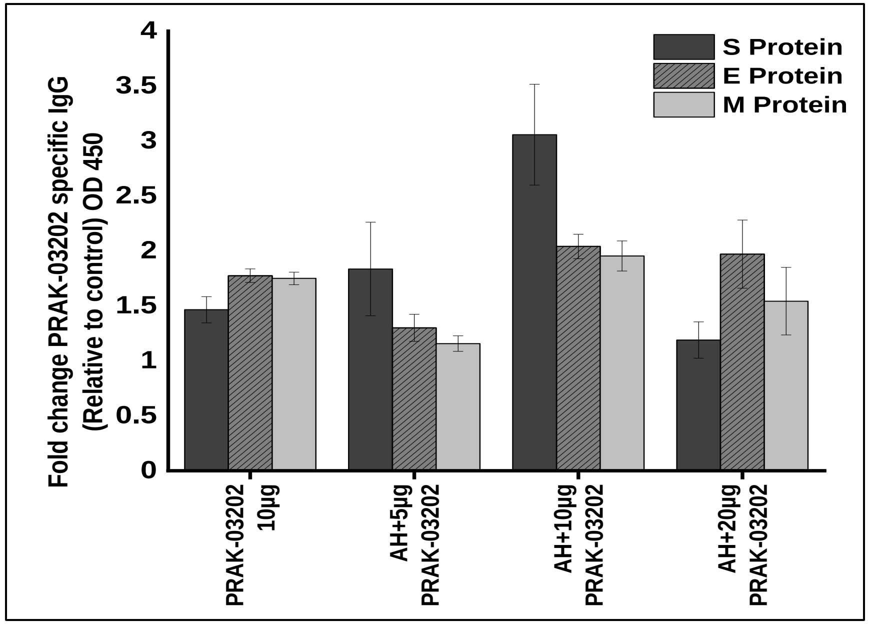

Figure 4A

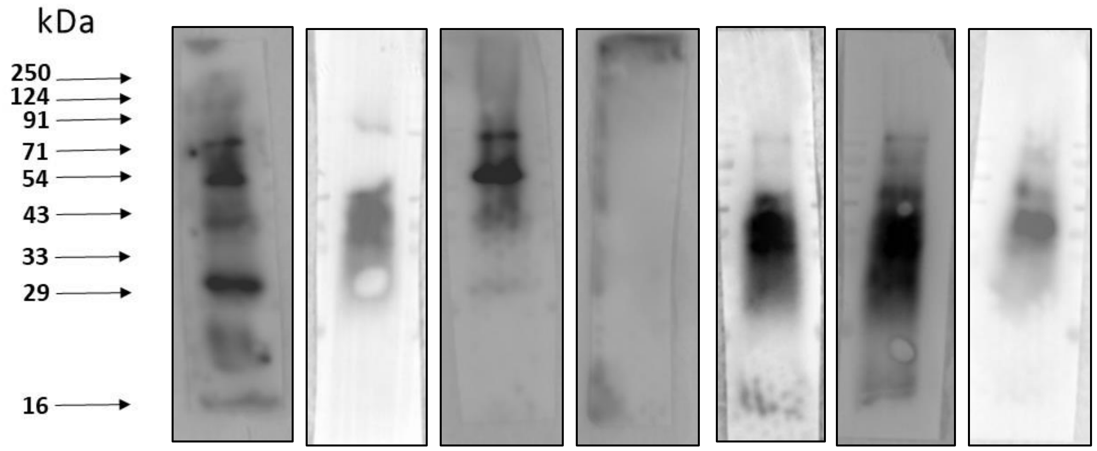


Figure 4B

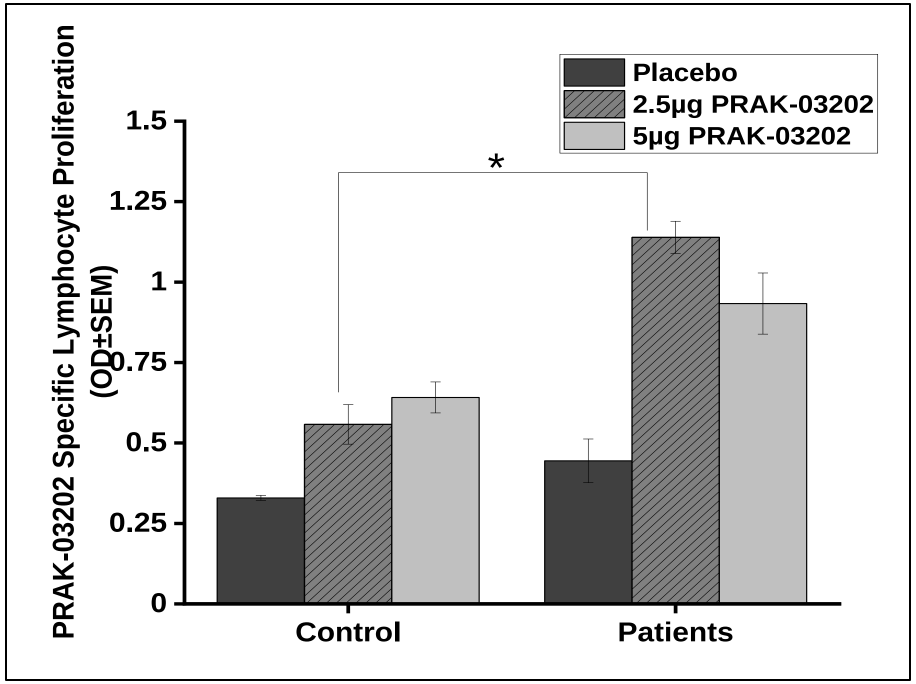


Figure 4C

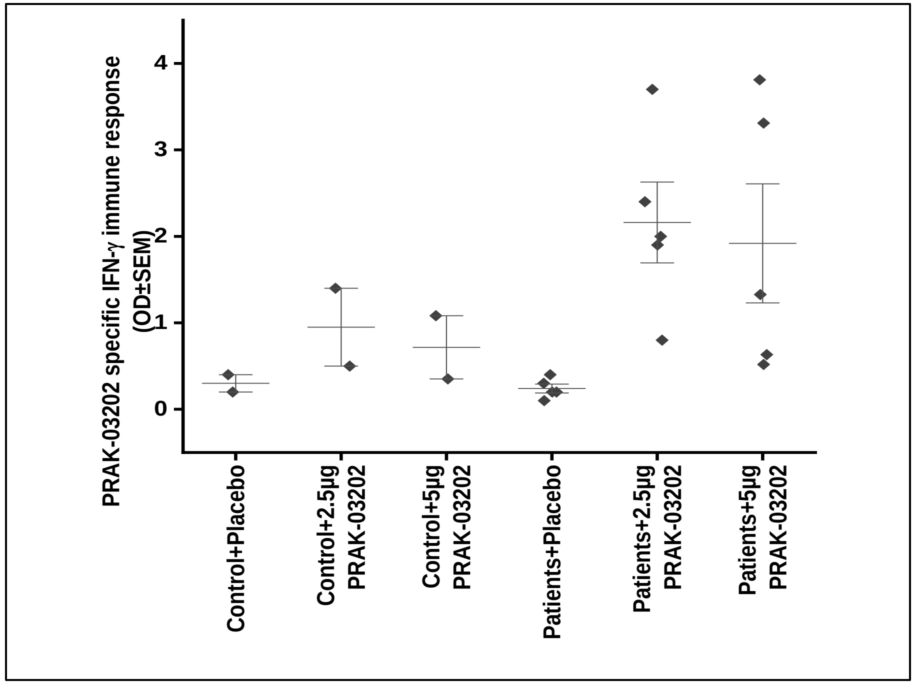

**Fig 5**

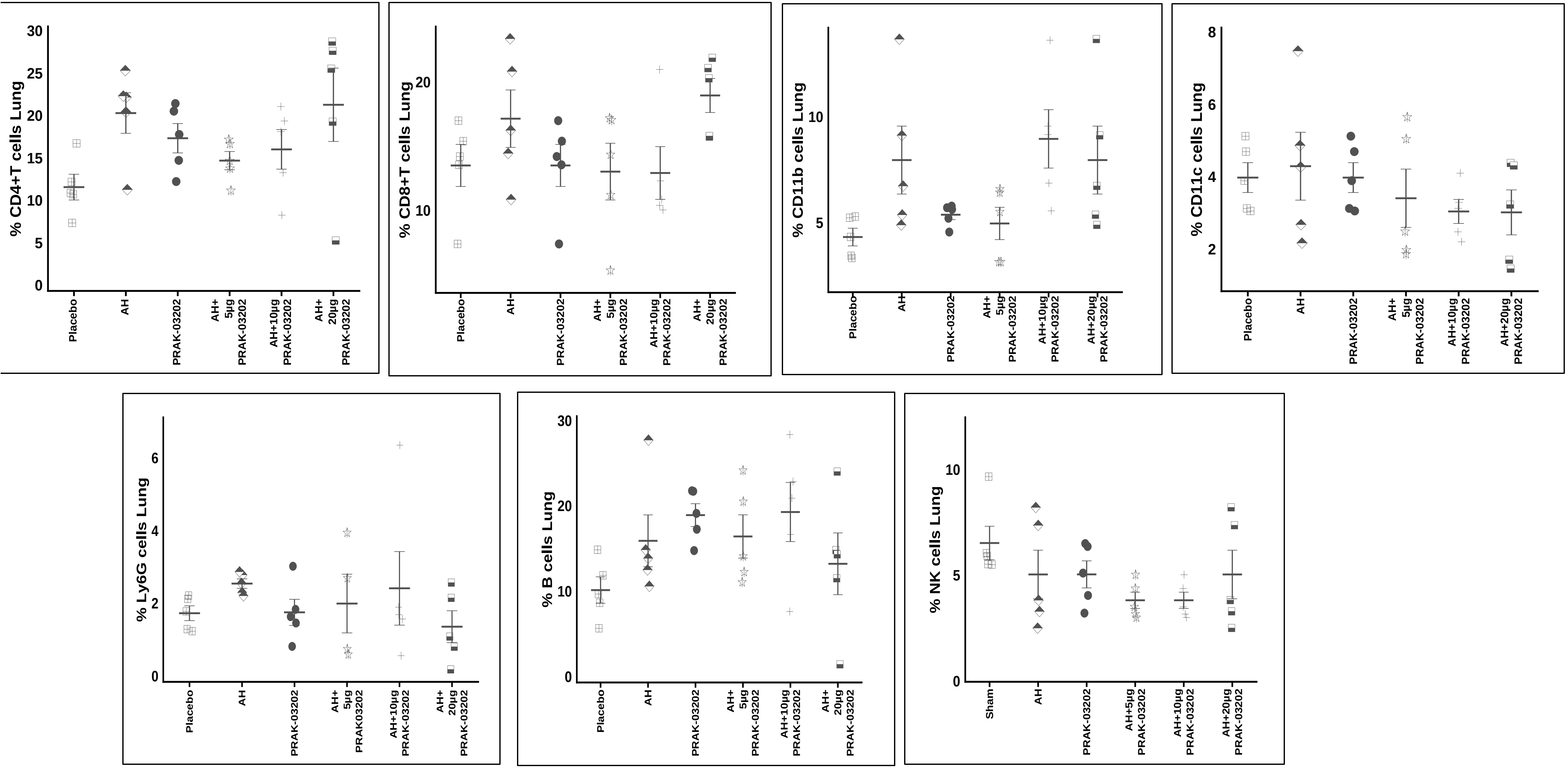
% CD4+T cells Lung
0
5
10
15
20
25
30
Placebo
AH
PRAK-03202
AH+ 5µg PRAK-03202
AH+10µg PRAK-03202
AH+ 20µg PRAK-03202
% CD8+T cells Lung
10
20
Placebo
AH
PRAK-03202
AH+ 5µg PRAK-03202
AH+10µg PRAK-03202
AH+ 20µg PRAK-03202
% CD11b cells Lung
5
10
Placebo
AH
PRAK-03202
AH+ 5µg PRAK-03202
AH+10µg PRAK-03202
AH+20µg PRAK-03202
% CD11c cells Lung
2
4
6
8
Placebo
AH
PRAK-03202
AH+ 5µg PRAK-03202
AH+10µg PRAK-03202
AH+20µg PRAK-03202
% Ly6G cells Lung
0
2
4
6
Placebo
AH
PRAK-03202
AH+ 5µg PRAK03202
AH+10µg PRAK-03202
AH+ 20µg PRAK-03202
% B cells Lung
0
10
20
30
Placebo
AH
PRAK-03202
AH+ 5µg PRAK-03202
AH+10µg PRAK-03202
AH+ 20µg PRAK-03202
% NK cells Lung
0
5
10
Sham
AH
PRAK-03202
AH+5µg PRAK-03202
AH+10µg PRAK-03202
AH+20µg PRAK-03202

Fig 6 A

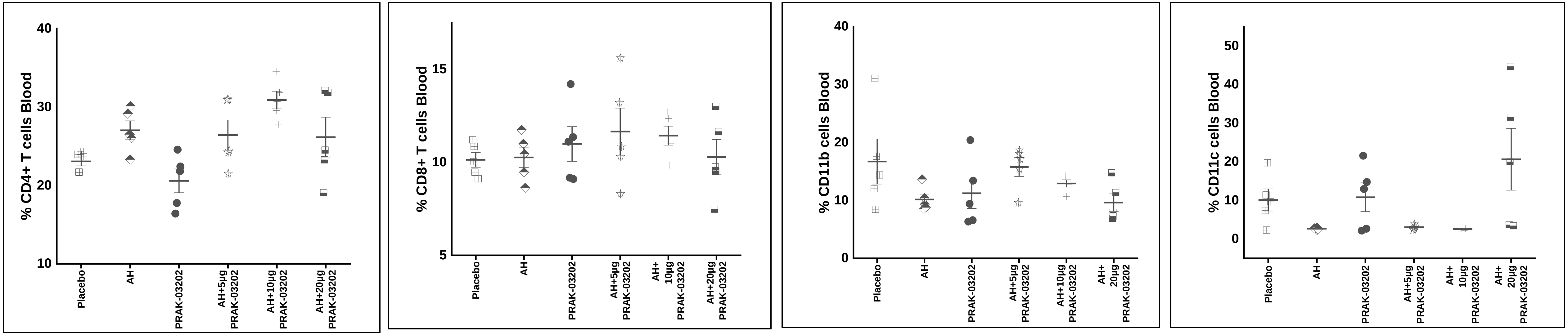
% CD4+ T cells Blood
% CD8+ T cells Blood
% CD11b cells Blood
% CD11c cells Blood
Placebo
AH
PRAK-03202
AH+5µg PRAK-03202
AH+10µg PRAK-03202
AH+20µg PRAK-03202

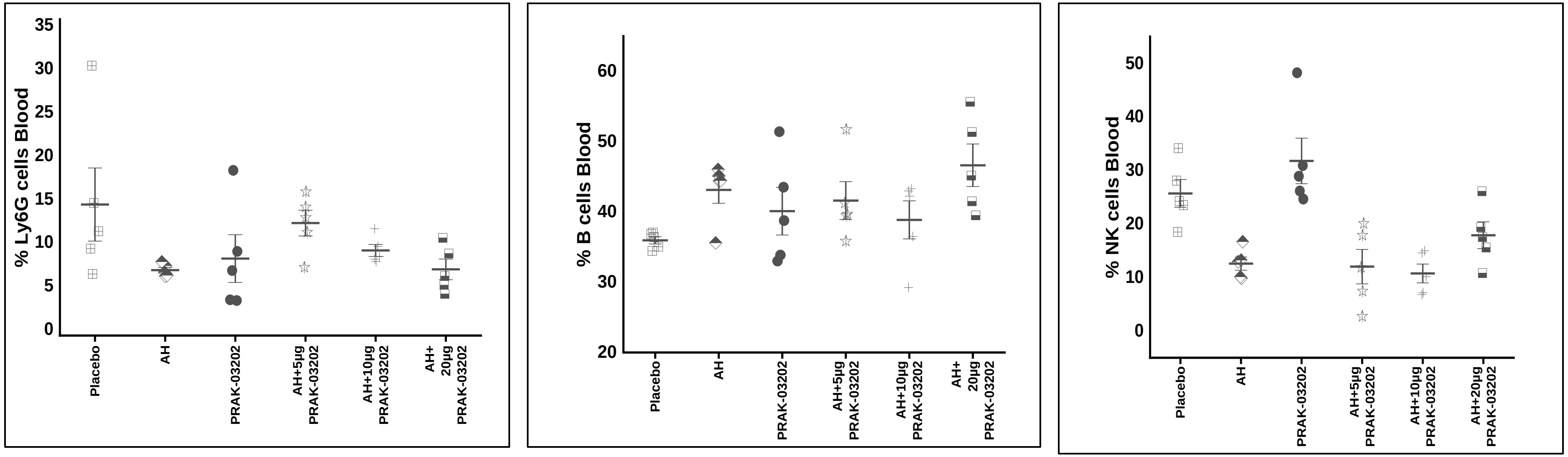
% Ly6G cells Blood
% B cells Blood
% NK cells Blood
Placebo
AH
PRAK-03202
AH+5µg PRAK-03202
AH+10µg PRAK-03202
AH+ 20µg PRAK-03202

Fig 6 B

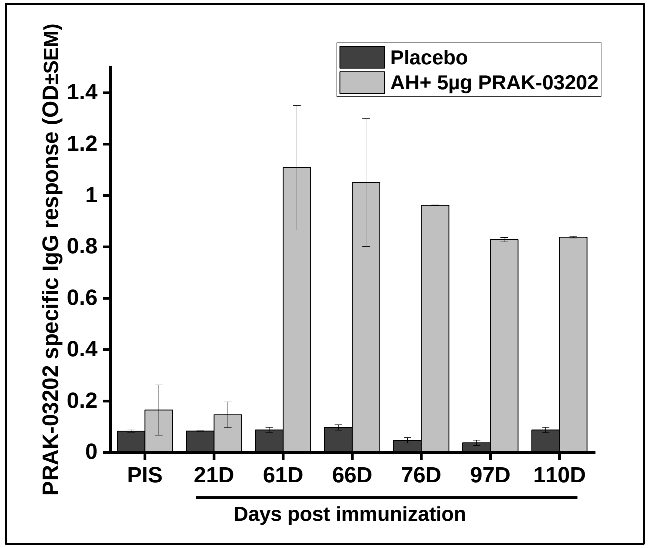
Placebo
AH+ 5µg PRAK-03202
PRAK-03202 specific IgG response (OD±SEM)
PIS
21D
61D
66D
76D
97D
110D
Days post immunization
